# Zika virus spreads through infection of lymph node-resident macrophages

**DOI:** 10.1101/2022.09.20.508424

**Authors:** Glennys V. Reynoso, David N. Gordon, Anurag Kalia, Cynthia C. Aguilar, Courtney S. Malo, Maya Aleshnick, Kimberly A. Dowd, Christian R. Cherry, John P. Shannon, Sophia M. Vrba, Sonia Maciejewski, Kenichi Asano, Theodore C. Pierson, Heather D. Hickman

## Abstract

Zika virus (ZIKV) is an arthopod-vectored flavivirus that disseminates from the infection site into peripheral tissues, where it can elicit virus-induced pathology. To move through the body, ZIKV is thought to exploit the mobility of myeloid cells, in particular monocytes and dendritic cells. However, multiple distinct steps during viral spread culminate in peripheral tissue infection, and the timing and mechanisms underlying mobile immune cell shuttling of virus remain unclear. To understand the very early steps in ZIKV dissemination from the skin, we kinetically and spatially mapped ZIKV-infected lymph nodes (LNs), an intermediary stop *en route* to the blood. Contrary to dogma, migratory immune cells were not required for large quantities of virus to reach the LN or blood. Instead, ZIKV rapidly infected a subset of immobile macrophages in the LN, which shed virus through the lymphatic pathway into the blood. Importantly, infection of LN macrophages alone was sufficient to initiate viremia. Together, our studies indicate that sessile macrophages that live and die in the LN contribute to initial ZIKV spread to the blood. These data build a more complete picture of ZIKV movement through the body and identify an alternate anatomical site for potential antiviral intervention.

**Highlights:**

- ZIKV infects and replicates in distinct LN macrophage populations
- LN macrophage infection results in infectious virus in the blood
- Virus reaches the blood in the absence of DC migration or monocyte infection
- Nodal macrophage infection does not sustain viremia or induce morbidity

**Graphical abstract:** Caption:
ZIKV infects lymph node macrophages, which shed infectious virus into the lymph and then blood.

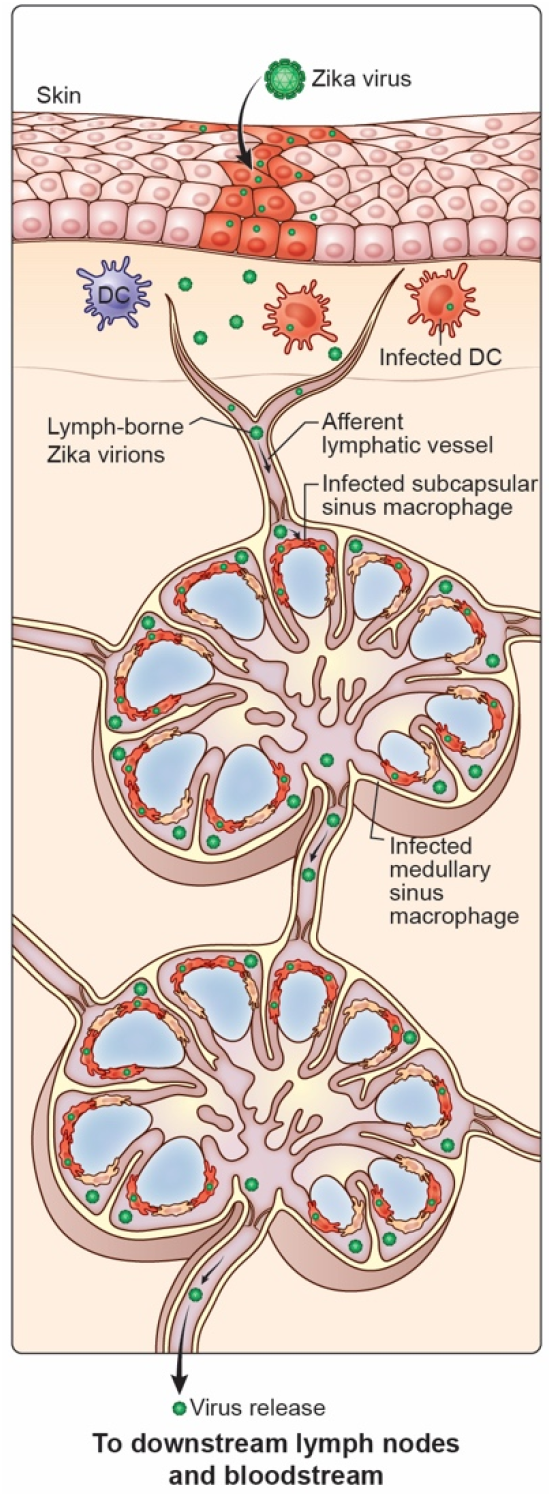

## Introduction

Flaviviruses are single-stranded RNA viruses that cause a spectrum of severe neurotropic and viscerotropic human diseases and have high epidemic potential (Pierson and Diamond, 2020). Zika virus (ZIKV) is a mosquito-borne flavivirus that explosively emerged in the Americas in 2015 (Diamond et al., 2019). Although most ZIKV infections reported during this epidemic were mild and self-limiting, some individuals experienced prolonged viremia with viral RNA detectable in the serum for weeks to months (Paz-Bailey et al., 2018). This epidemic revealed new clinical features of infection, including a range of congenital neurodevelopmental diseases, such as microcephaly (Mlakar et al., 2016). Concerns over ZIKV-induced disease spurred the rapid development of new animal models in which to evaluate ZIKV pathogenesis, vaccines candidates, and therapeutics (reviewed in (Shan et al., 2018)). Neutralizing Abs (NAbs) have been identified as a correlate of protection after viral challenge (Abbink et al., 2017; Dowd et al., 2016b; Richner et al., 2017; Van Rompay et al., 2019). No drugs or vaccines are currently approved for treatment of infection.

Zika virus induces pathogenesis after dissemination from the skin into the blood, after which the virus gains access to many different peripheral organs and tissues, including the brain and, in pregnant women, the fetus(Gorman et al., 2018; Lazear et al., 2016; Morrison and Diamond, 2017). Both viral replication in the tissue as well as the induction of antiviral immune responses, including type I interferons (IFN-Is), can cause tissue damage and fetal demise(Miner et al., 2016; Yockey et al., 2018). The importance of dissemination in ZIKV-induced disease has led to intensive investigation of the cellular targets for ZIKV infection and the routes of dissemination into peripheral tissues. The TAM family of receptor tyrosine kinases (including Tyro3, Axl, and Mer) were among the first proteins identified that could promote ZIKV infection *in vitro* (Liu et al., 2016; Mladinich et al., 2017; Richard et al., 2017). As these proteins are highly expressed on endothelial cells(Lemke, 2013), ZIKV infection of vascular endothelial cells via TAM receptors provides a simple explanation for the ability of ZIKV to access many different tissues from the blood. However, mice deficient in Axl and Mertk were soon shown to exhibit similar levels of ZIKV infection and viral distribution as wild-type mice(Hastings et al., 2017). A second attractive route for ZIKV movement into the tissues is along with mobile immune cells. In human blood, circulating monocytes were identified as the primary cell type infected by ZIKV, and human monocytes can also be productively infected in vitro (Ayala-Nunez et al., 2019; Foo et al., 2017; Michlmayr et al., 2017). Using myeloid-restricted ZIKV to infect mice, McDonald *et al*. concluded that monocytes represented the major myeloid population to disseminate ZIKV(McDonald et al., 2020). Indeed, monocytes represent an attractive conduit for viral movement, as these cells are abundant in the blood and readily migrate into even immune-privileged tissues(Teh et al., 2019). Thus, monocytes have been suggested to serve as a “Trojan horse” that distributes ZIKV throughout the body(Ayala-Nunez *et al*., 2019; Jurado and Iwasaki, 2017). Many other myeloid cells in the skin, including dermal dendritic cells and Langerhans cells, also become infected by ZIKV(Cerny et al., 2014; Duangkhae et al., 2018; Hamel et al., 2015; McDonald *et al*., 2020; Schmid and Harris, 2014; Wu et al., 2000) and have been proposed to facilitate ZIKV dissemination.

Viral dissemination is often viewed as the sum of its products (e.g., peripheral organ infection), as this is ultimately the step that results in virus-induced disease. However, dissemination is a complex, multi-step process and the disruption of very early dissemination events could potentially eliminate downstream viral entree into the tissue. However, the cells responsible for early viral movement, particularly within the lymph node (LN) are unclear. Additionally, it is unknown whether the LN serves to amplify virus draining from the infection site or simply as a passageway for virus produced in the skin.

In this study, we examined the early events after ZIKV infection to uncover the mechanisms allowing the initial movement of ZIKV from skin to LN to blood. We provide evidence that mobile immune cells do not initially act as Trojan horses that accidentally distribute ZIKV while trying to perform their immune functions. Instead, we show that LNs are seeded by infectious ZIKV within minutes of inoculation via lymphatic vessel transport (rather than cellular migration). Importantly, immobile macrophages in the LN are the primary source of infectious virus that first reaches the bloodstream. This viremia can be established without monocyte infection or dendritic cell migration. Thus, LN macrophages are link in the chain of ZIKV dissemination that could be targeted by antiviral therapeutics.

## Results

### ZIKV is captured by the local draining lymph node before dissemination

To understand how ZIKV disseminates from the skin, we used a common mouse model of ZIKV infection, footpad inoculation of *Ifnar1*^-/-^ mice (Morrison and Diamond, 2017). Although ZIKV efficiently antagonizes human type I interferon (IFN-I) responses, it cannot bind to murine STAT2, necessitating circumvention of IFN-I signaling to support murine infection (Grant et al., 2016). We first examined the very early kinetics of viral delivery to the LN draining the hind foot, the popliteal (PLN), reasoning that we should not detect high levels of virus within an hour (hr) if skin cells were needed to replicate virus (**Figure 1**). We infected mice in the hind footpads with 10^4^ focus-forming units (FFU) of ZIKV H/PF/2013 and harvested PLNs from 5 minutes (min) to 1 hr post-infection (p.i.) for viral titers as determined by a focus-forming assay (FFA) (**Figure 1A**). Within the first hr, we detected infectious ZIKV in PLN homogenates, but not in the serum, indicating that virus reaches the PLN after initial inoculation via the lymphatics. The initial entry of infectious virus in the PLN peaked between 10 and 30 min after inoculation before dropping to levels just above the limit of detection at 1 hr.

**Figure 1:**
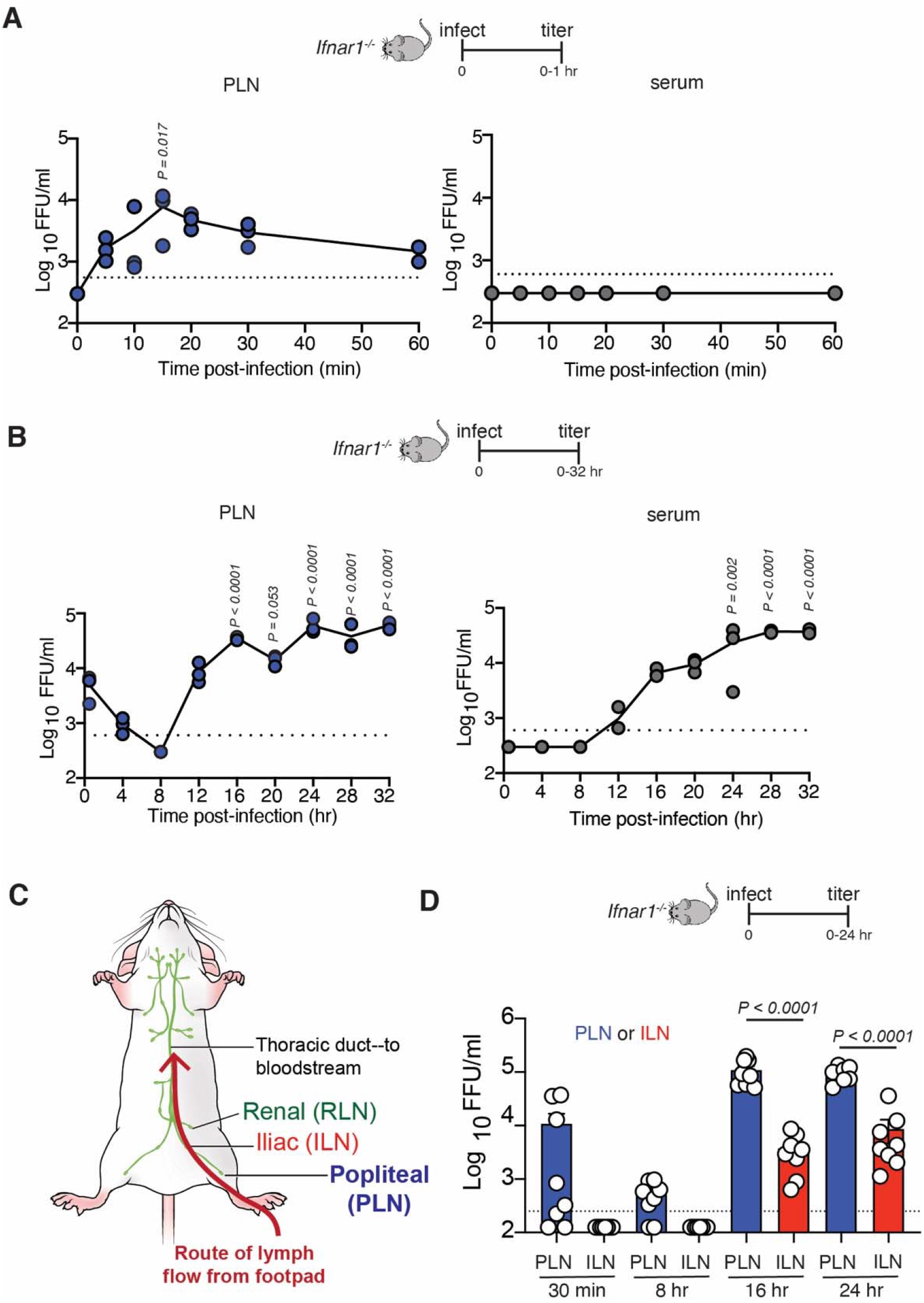
ZIKV replicates in the PLN before systemic dissemination. **(A)** Viral titers (in focus-forming units (FFU)/ml)) in the PLN (left, blue dots) or serum (right, grey dots) harvested at the indicated time (in min) during the first hr following footpad (FP) injection of *Ifnar1*^-/-^ mice with 10^4^ FFU of ZIKV H/PF/2013. Dots represent individual mice (either pooled PLNs or separate sera) and the average of technical replicates in the focus-forming assay (FFA). A dashed line shows the limit of detection of the FFA. Values below half the LOD are reported as half the LOD (125 FFU/ml). PLNs and sera were harvested from the same mice. Data are shown from 1 of 2 complete timecourse experiments. **(B)** Viral titers in the PLN (left, blue dots) or serum (right, grey dots) at the indicated timepoint during the first 32 hr p.i. of *Ifnar1*^-/-^ mice. Data are shown from 1 of 2 complete timecourse experiments. **(C)** Schematic illustration of the route of lymphatic draining from the footpad. Lymph first flows to the popliteal LN (PLN, the “outpost” LN) followed by the iliac LN (ILN). Downstream from nodes, lymph enters the thoracic duct followed by the subclavian vein (where it enters the blood). **(D)** Viral titers in PLNs (blue bars) or ILNs (red bars) of *Ifnar1*^-/-^ mice at the indicated time p.i. (in hr). PLNs and ILNs were harvested from the same mice. Data are shown from two pooled experiments with four mice/group. All experiments were repeated 2-3 times with 3-4 mice per group. Dots represent individual mice (either pooled LNs or separate sera) and the average of technical replicates in the FFA. Error bars = SEM. Dashed line = limit of detection for assay. Statistics =one-way ANOVA. Exact *P* values are shown in relation to uninfected controls.

We next quantified infectious virus in the PLN or serum over a longer period--every 4 hr for the first 32 hr p.i. (**Figure 1B**). At 8 hr, we did not detect infectious virus in the PLN; however, viral titers rebounded to greater levels than the initial input virus by 16 hr p.i. Viremia was detected at 12 hr p.i. and became elevated by 16 hr p.i. We obtained similar results in wild-type C57BL/6 mice receiving the anti-IFNAR1 Ab MAR1-5A3 (MAR1, described in (Lazear *et al*., 2016)) (**Supplementary Figure 1**). In C57BL/6 mice with intact IFN-I signaling, similar levels of infectious virus arrived at the PLN, but titers never increased, indicating that viral replication drives the higher viral titers seen in the PLN at later timepoints in *Ifnar1^-/-^* mice (**Supplementary Figure 1**).

The LNs act as a system of sequential filters that remove viruses from the lymph flow (Ozawa et al., 2022). From the PLN, lymph fluid passes to the iliac LN and then to the renal LN before eventually emptying into the bloodstream (schematically depicted in **Figure 1C**) (Harrell et al., 2008). Thus, the iliac should have the opportunity to sequester any virus not captured by the PLN. Viral titers in the iliac LN detected no infectious virus before 16 hr p.i., indicating that the PLN captures all the lymph-borne virus at this initial inoculating dose (**Figure 1D**). Together, these data demonstrate that infectious ZIKV inoculated into the skin passes to the draining LN, where it is captured.

### ZIKV replicates in distinct lymph node macrophage niches

Lymph nodes contain at least five distinct populations of macrophages, which form a network to carry out specific immune functions (Bellomo et al., 2018). Macrophages present in nodal sinuses represent an immobile population of phagocytic cells that are situated with their bodies in the sinusoidal space and projections through lymphatic endothelial cells that anchor them to the sinus floor while providing cytokines necessary for their survival (Gray and Cyster, 2012; Mondor et al., 2019). Sinus-resident macrophages can be further sub-divided based on their location within the LN and expression of cell-surface receptors (Phan et al., 2009). Sinus-resident macrophages nearest the afferent (incoming) lymph vessel have been termed subcapsular sinus macrophages (SSMs) while those nearest the efferent (exiting) lymph vessels are known as medullary sinus macrophages (MSMs). These populations access lymph-borne particulates sequentially as lymph enters the node through the subcapsular sinus and flow around to the medullary sinus (Jafarnejad et al., 2015).

Because of their location, we first investigated whether lymphoid sinus-resident macrophages captured ZIKV. We first imaged frozen cross-sections of the PLN harvested from 8-24 hr p.i. from C57BL/6, C57BL/6 treated with MAR1, or *Ifnar1*^-/-^ mice (**Figure 2**). LNs were carefully oriented such that they were sectioned from the top of the LN (near the afferent lymphatics) through the medulla and hilum as previously described (Reynoso et al., 2019), thus affording a complete view of the major anatomical subdivisions of the LN. We distinguished CD169^+^sinusoidal macrophage populations by their location and morphology as previously described (Phan *et al*., 2009; Reynoso *et al*., 2019; Sung et al., 2012). At 8 hr p.i., we detected ZIKV E protein that colocalized with small patches of SSMs, regardless of the type I IFN-signaling capacity of the recipient mice and thus likely reflecting binding of input virus (**Figure 2A**). We did not detect E protein staining in control LNs infected with a different virus to induce nodal inflammation, which increases background Ab staining (vaccinia virus (VACV)) (**Supplementary Figure 2**). We identified bonafide ZIKV-infected cells through staining for the ZIKV non-structural protein NS2b, which is not incorporated into virions (**Figure 2B-D**). By 8 hr p.i., NS2b staining was detected at low levels in the LN and was restricted to SSMs. By 16 hr, we detected prominent NS2b staining in MSMs located in medullary sinuses (**Figure 2C and D**). Most of the NS2b-infected cells express CD11b and SIGNR1, but not CD11c at 16 hr p.i. (consistent with our previous analyses demonstrating a paucity of infected DCs in ZIKV-infected LNs at early timepoints (Reynoso *et al*., 2019) (**Figure 2E-G**). Staining of the ILN revealed a similar but delayed pattern of infection, with SSMs being productively infected at 16 hr p.i. followed by MSMs at 24 hr p.i. (**Supplementary Figure 2**). Together with Figure 1, these data reveal that infection in *Ifnar1*^-/-^ mice proceeds through macrophage networks along the route of lymph flow.

**Figure 2:**
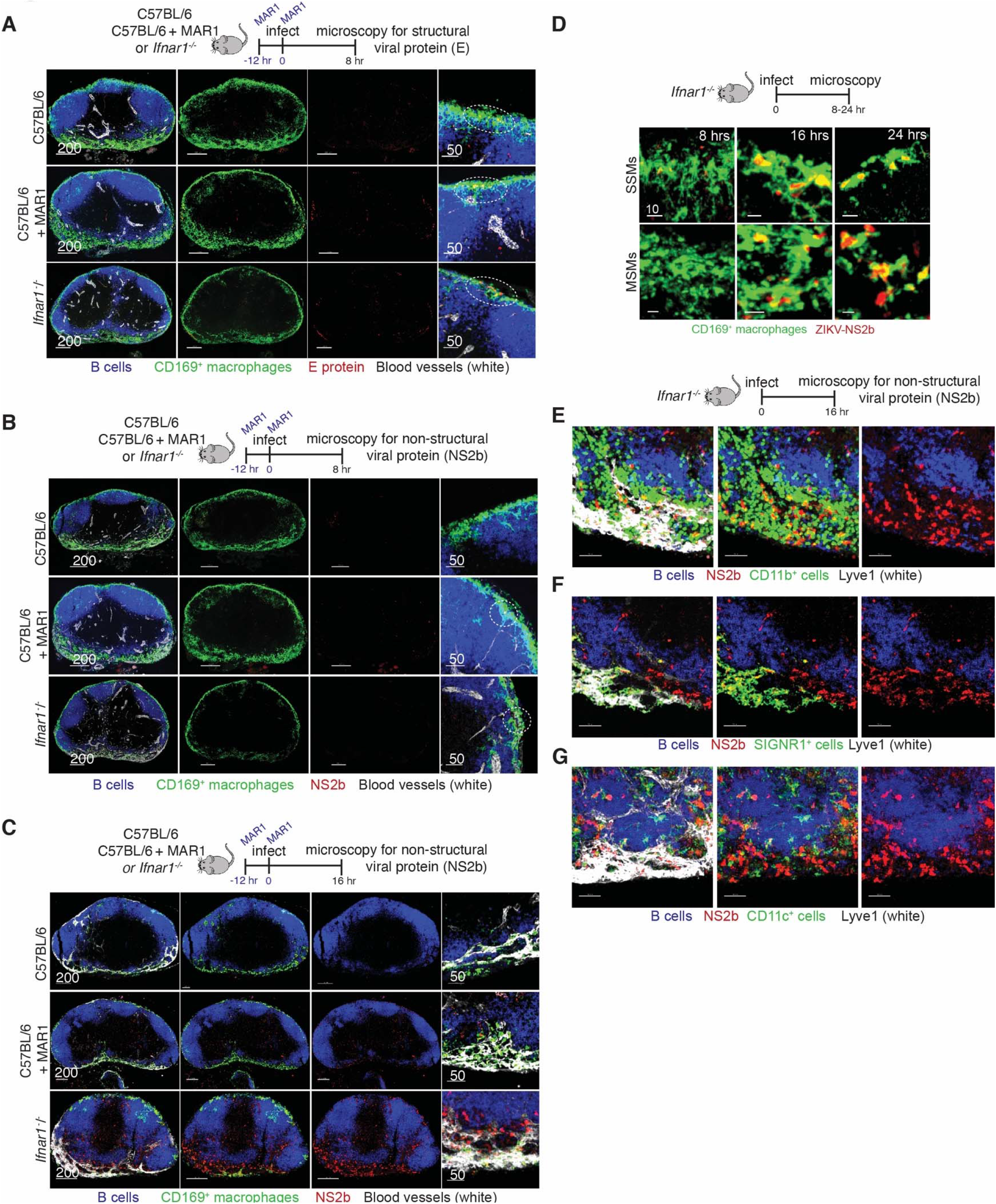
ZIKV infects macrophages in lymph node sinuses. **(A)** Confocal images of frozen PLN sections from nodes harvested 8 hr p.i. from C57BL/6 (top panels), C57BL/6 mice given the a-IFNAR1 Ab MAR1 (middle panels), or *Ifnar1*^-/-^ (bottom panels) mice. Blue = B cells; green = CD169^+^ macrophages; red = ZIKV E protein; white = CD31 (blood vessels). Far right panel shows a higher magnification view of the subcapsular sinus. Dashed ovals show specific areas with ZIKV E protein staining. **(B)** Confocal images of frozen PLN sections from nodes harvested 8 hr p.i. from C57BL/6 (top panels), C57BL/6 mice given the a-IFNAR1 Ab MAR1 (middle panels), or *Ifnar1*^-/-^ (bottom panels) mice. Blue = B cells; green = CD169^+^ macrophages; red = ZIKV NS2b protein; white = CD31 (blood vessels). Far right panel shows a higher magnification view of the subcapsular sinus. Dashed ovals show specific areas with ZIKV NS2b protein staining. **(C)** As in (b) but PLNs were harvested 16 hr p.i. **(D)** Confocal images of frozen PLN sections from nodes harvested at the indicated time (shown in hr p.i. in the upper right corners) from *Ifnar1*^-/-^ mice. Top panels show macrophages in the subcapsular sinus (SCS macrophages, SSMs) and bottom panels show macrophages in medullary sinuses (MS macrophages, MSMs). Green = CD169^+^ macrophages; red = ZIKV NS2b protein. **(E-G)** Confocal images of frozen LN sections from nodes harvested 16 hr p.i. from *Ifnar1*^-/-^ mice. White = Lyve1; red = ZIKV NS2b protein. Green = CD11b (**E)**; SIGNR1 (**F**); or CD11c (**G**). Images are representative of 6-10 PLNs/timepoint/condition harvested from 3-5 mice. Scalebars =μms.

### ZIKV infection disrupts lymph node macrophages

Viral drainage leads to the death or attrition of SSMs, which can impair the development of antiviral humoral responses (Gaya et al., 2015; Sagoo et al., 2016). We exploited this feature as a surrogate measure of LN-macrophage sensing of lymph-borne ZIKV (independent of direct macrophage infection). We performed flow cytometry on single-cell suspensions of popliteal or iliac LNs harvested 72 hr p.i. from mice with or without IFN-I signaling (**Figure 3A and B**). We gated on live CD45^+^ CD11c^lo^ CD11b^+^ CD169^+^ cells and further separated macrophage populations using F4/80 (present on MSMs only (Phan *et al*., 2009)). SSMs were absent in both the PLN and iliac LNs at this timepoint; this did not require productive ZIKV infection as SSMs were also ablated in WT mice. In contrast to SSMs, MSMs have not been shown to undergo post-infection attrition (Gaya *et al*., 2015). However, by 72 hr. p.i., MSMs were largely absent in *Ifnar1*^-/-^ mice, although some MSMs were still detectable in C57BL/6 mice with intact IFN-I signaling (**Figure 3**). The pronounced MSM ablation in *Ifnar1*^-/-^ mice indicates that the MSM compartment has access to virus or viral Ag produced after viral replication in these animals.

**Figure 3:**
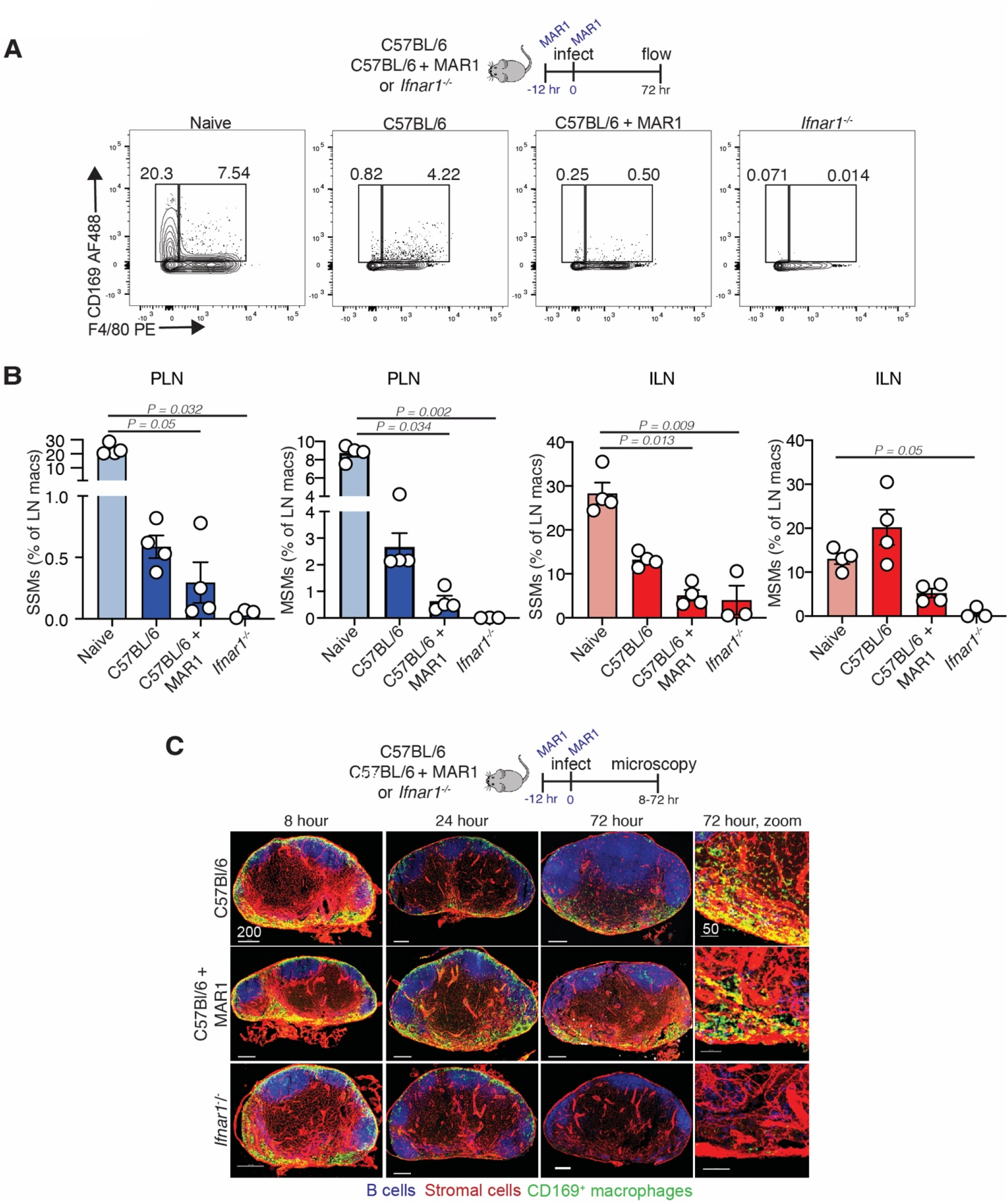
Zika virus disrupts LN macrophage networks. **(A)** Flow plots generated from single cell suspensions of PLNs harvested from uninfected C57BL/6 (left panel), or from ZIKV-infected C57BL/6 (middle left), C57BL/6 + MAR1 Ab (middle right) or *Ifnar1*^-/-^ (right) mice 72 hr p.i. Cells were first gated on CD45^+^ B220^-^ CD3_-_ CD11b^+^CD11c^low^ cells. Gating indicates SSMs (left gates, CD169^+^ F4/80^-^) and MSMs (right gates, CD169^+^ F4/80^+^). **(B)** Frequency of CD169^+^ SSMs (far left and middle right panels) or MSMs (middle left and far right panels) in the either PLNs (blue bars) or ILNs (red bars) as a percentage of total LN macrophages from experiment shown in flow plots in (A). Statistics = One-way ANOVA. Dots show pooled LNs from individual mice. Scalebars = SEM. Experiment was repeated 3 times with 3-4 mice/group. **(C)** Confocal images of frozen PLN sections harvested from C57BL/6 (top panels), C57BL/6 mice given the a-Ifnar1 Ab MAR1 (middle panels), or *Ifnar1*^-/-^ (bottom panels) mice at the indicated time p.i. Blue = B cells; green = CD169^+^ macrophages; red = stromal cells (ERTR-7^+^). Far right panel shows a higher magnification view of the medullary sinus. Images in **C** are representative of 6-10 LNs/timepoint/condition harvested from 3-5 mice. Scalebars =μms.

LN macrophages are difficult to remove by enzymatic tissue dissociation and may be overestimated by *ex vivo* approaches (Gray et al., 2012), therefore, we also examined LNs over the same time frame using confocal microscopy (**Figure 3C**). Imaging at 72 hr p.i. recapitulated the SSM and MSM ablation in *Ifnar1*^-/-^ mice observed using flow. We also noted little overt change in LN architecture (evaluated through the staining of LN stromal and vascular cells). Interestingly, the LN macrophage network remained largely intact at 8 hr and partially intact at 24 hr (**Figure 3C**), thus suggesting that early disruption and failure to capture virus draining from the skin was not likely to account for the appearance of virus in the blood at 12 hr p.i.. Together, these data show that skin infection with ZIKV results in the post-dissemination disruption of the LN macrophage network.

### LN macrophage infection alone allows systemic viral dissemination

To more conclusively understand the role of LN macrophages in preventing or allowing ZIKV dissemination, we crossed *Siglec1-cre* mice (with Cre recombinase expression in CD169^+^ cells (Karasawa et al., 2015)) with *Ifnar1-floxed* mice (Prigge et al., 2015) to conditionally delete IFN-I signaling in CD169^+^ LN macrophages (*Siglec1-cre Ifnar1^fl/fl^* mice, hereafter referred to as CD169 cKO, **Figure 4**). Viral titers in the popliteal or iliac LNs at 24 hr p.i. infection were surprisingly high even though viral replication was restricted to only LN macrophages (titers were statistically similar to *Ifnar1*^-/-^ mice) (**Figure 4A**). More importantly, infectious ZIKV in the serum was similar in mice with unrestricted viral replication as in CD169 cKO mice, indicating that infection of LN macrophages alone could lead to high levels of disseminated virus by 24 hr p.i. Confocal analyses confirmed LN macrophage replication (via ZIKV NS2b protein staining) in both the popliteal and iliac LNs of CD169 cKO mice (**Figure 4B and C**). These data indicate that, after ZIKV infection, the sinusoidal LN macrophage system is incapable of fully sequestering virus but instead releases virus to initiate systemic infection.

**Figure 4:**
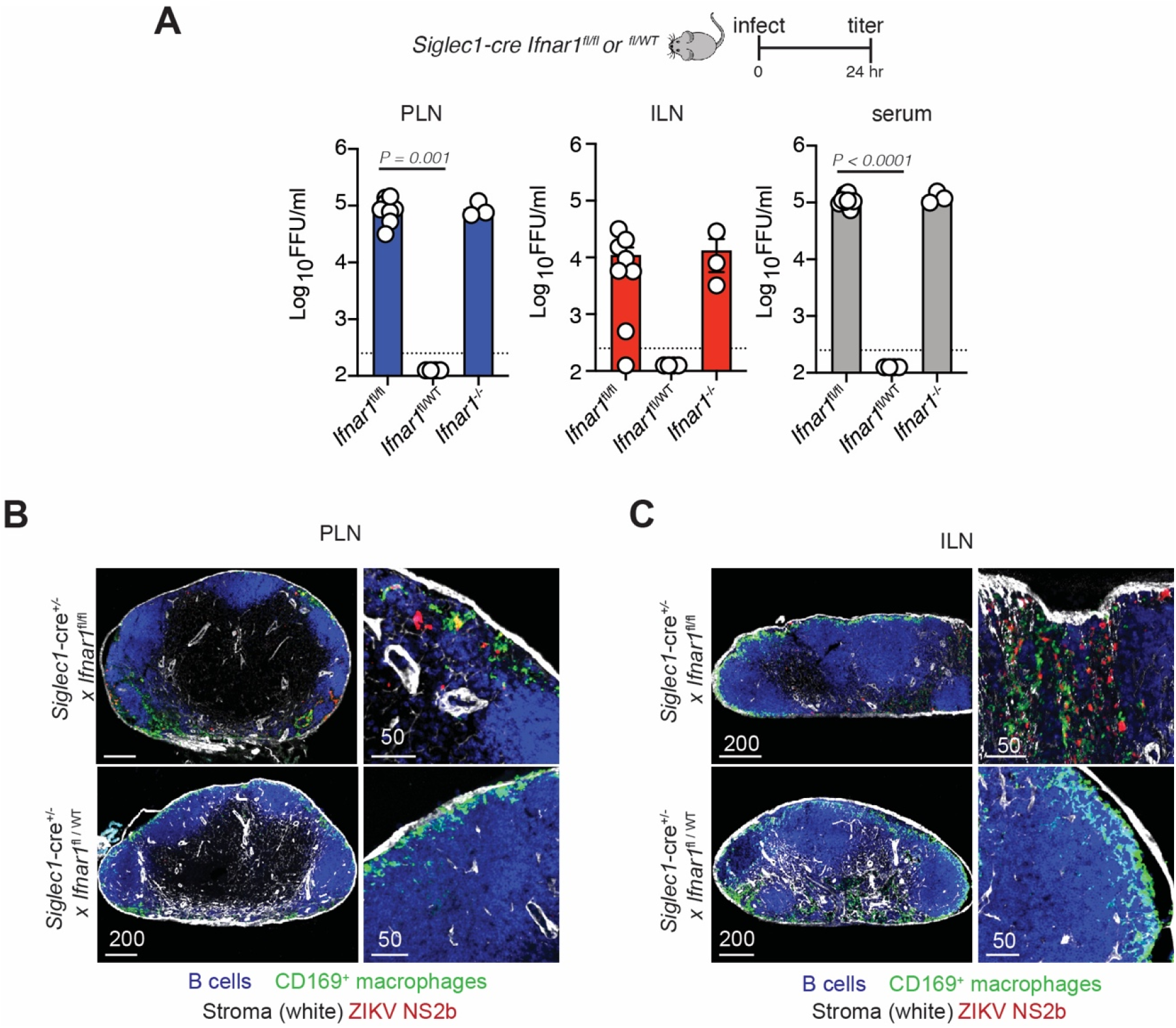
Lymph node macrophage infection alone supports systemic dissemination. **(A)** Viral titers (FFU/ml) in the PLN (left, blue bars), ILN (middle, red bars), or serum (right, grey bars) harvested 24 hr p.i. of *Siglec1-cre Ifnar*^fl/fl^ (homozygous *Ifnar1* knockout in CD169^+^ cells (cKO)) mice, *Siglec1-cre Ifnar*^fl/WT^ (heterozygous knockout in CD169^+^ cells) or *Ifnar1*^-/-^ mice with 10^4^ FFU of ZIKV. Experiment was repeated 3 times with 3-4 mice per group. Results shown are pooled from two independent experiments. Dots represent individual mice (either pooled LNs or separate serum) and the average of technical replicates (values of zero are not shown due to log scale). Dashed line = limit of detection for assay. Values below half the LOD are reported as half the LOD (125 FFU/ml). **(B)** Confocal images of frozen PLN sections harvested 24 hr p.i. from *Siglec1-cre Ifnar*^fl/fl^ (homozygous cKO) mice (top panels) or *Siglec1-cre Ifnar*^fl/WT^ (heterozygous cKO mice, bottom panels) stained for B cells (using B220 Ab, (blue)); CD169^+^ macrophages, (green); stromal cells (using ERTR7 (white)); and ZIKV NS2b protein (red). Right panels shower higher magnification images. Scalebars =μms. **(C)** As in (B) but showing ILN instead of PLN.

### Dermal DC migration is not required for viral movement to the blood

Although dermal DCs are thought to carry infectious virus to the LN to initiate nodal infection and adaptive immune responses, our kinetic examination of nodal macrophage infection suggested that macrophages were infected before DC trafficking from the skin, which is estimated to require approximately 12 hr. Therefore, we next quantitated virus at the infection site (foot), LN and serum of CD169 cKO mice at 12 hr p.i. (**Figure 5A**). Although we detected viral dissemination in the serum by 12 hr in both *Ifnar1*^-/-^ and CD169 cKO mice, we did not yet detect infectious virus in foot homogenates (**Figure 5A**). Viral titers were statistically similar between complete Ifnar1 knockouts and CD169 cKO mice at this early time point.

**Figure 5:**
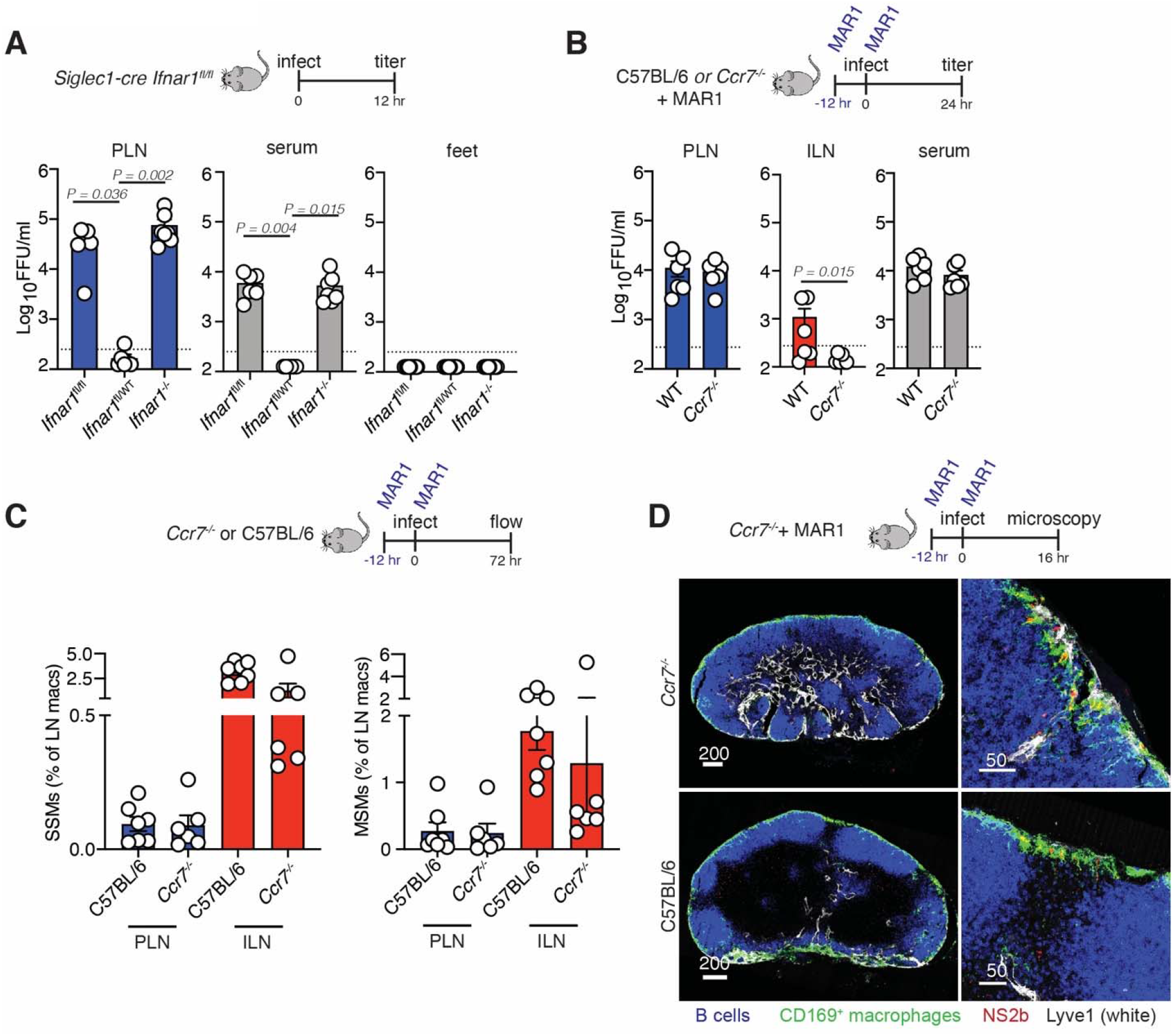
Migrating dendritic cells are dispensable for ZIKV dissemination. **(A)** Viral titers (FFU/ml) in the PLN (left, blue bars), serum (middle, grey bars), or footpad (right, yellow bars) harvested 12 hr p.i. from *Siglec1-cre Ifnar*^fl/fl^ (homozygous *Ifnar1* knockout in CD169^+^ cells, cKO) mice, *Siglec1-cre Ifnar^fl/WT^* (heterozygous cKO mice) or *Ifnar1*^-/-^ mice with 10^4^ FFU of ZIKV. Experiment was repeated 2 times with 3 mice per group. Results shown are pooled from two independent experiments. Dots represent individual mice (either pooled LNs or separate serum or feet) and the average of technical replicates (values of zero are not shown due to log scale). Dashed line = limit of detection (LOD) for assay. Values below half the LOD are reported as half the LOD (125 FFU/ml). **(B)** Viral titers (FFU/ml) in the PLN (left, blue bars), ILN (middle, red bars), or serum (right, grey bars) harvested 12 hr p.i. from C57BL/6 or *Ccr7*^-/-^ mice treated with MAR1 and infected with 10^4^ FFU of ZIKV. Experiment was repeated 2 times with 3 mice per group. Results shown are pooled from two independent experiments. Dots represent individual mice (either pooled LNs or separate serum) and the average of technical replicates (values of zero are not shown due to log scale). Dashed line = limit of detection for assay. Values below half the LOD are reported as half the LOD (125 FFU/ml). **(C)** Frequency of CD169^+^ SSMs (left) or MSMs (right) in the either PLNs (blue bars) or ILNs (red bars) 72 hr p.i. of C57BL/6 or *Ccr7*^-/-^ mice as a percentage of total LN macrophages Statistics = One-way ANOVA. Dots show pooled LNs from individual mice. Scalebars = SEM. Experiment was repeated 2 times with 3 mice/group. **(D)** Confocal images of frozen LN sections from nodes harvested 16 hr p.i. from C57BL/6 or *Ccr7*^-/-^ mice treated with MAR1. Blue = B cells; green = CD169^+^ macrophages; red = ZIKV NS2b; white = Lyve1. Right panels show a higher magnification of infected SSMs. Scale bars =μms.

Skin-resident dermal DCs and Langerhans cells use the chemokine receptor CCR7 to migrate via the lymphatics to the skin-draining LN (Platt and Randolph, 2013). To dissect the contribution of DC/Langerhans migration from the skin to LN infection, we treated *Ccr7*^-/-^ mice (lacking DC lymphatic migration) with MAR1 and quantitated infectious virus at 24 hr p.i. (**Figure 5B**). Viral titers in the PLN were not statistically different regardless of DC migration and mice still developed similar levels of viremia. Notably, CCR7 deficiency significantly decreased viral titers in the ILN, suggesting cellular movement either from or within the PLN impacts viral transport to downstream LNs (as T cells also use CCR7 for inter- and intra-nodal migration (Braun et al., 2011)). Flow cytometry at 72 hr revealed attrition of SSMs and MSMs in both the popliteal and iliac LNs (**Figure 5C)**, indicating the DC migration is not needed for disruption of nodal macrophage networks. We confirmed productive infection of LN macrophages in MAR1-treated *Ccr7*^-/-^ mice through confocal microscopy and staining for NS2b (**Figure 5D**). Therefore, we conclude that LN macrophages can produce virus for systemic dissemination in the absence of a significant contribution of infectious virus from migratory DCs.

### Infected monocytes do not account for high levels of virus in the blood

Monocytes are mobile inflammatory myeloid cells rapidly recruited to sites of infection, where typically die or mature into sedentary tissue macrophages (Italiani and Boraschi, 2014). Flavivirus infection of both mice and humans results in monocyte mobilization from the bone marrow into the blood and then the skin, where they are thought to serve as targets of infection(Schmid and Harris, 2014) (Foo *et al*., 2017; Michlmayr *et al*., 2017). Further, monocytes have been proposed to become ZIKV “Trojan horses,” representing the major cell population disseminating virus in mice (McDonald *et al*., 2020). Based on these prior studies, we next assessed the role of classical Ly6c^high^ monocytes during early ZIKV dissemination (**Figure 6A and B**). Zika virus infection mobilized classical monocytes into the blood within 24 hr of infection. (**Figure 6B, left**). Although monocytes were detected in the infected PLN, treatment with MAR1 greatly reduced monocyte nodal numbers (**Figure 6B, right**). To deplete inflammatory monocytes, we treated mice with the depleting monoclonal Ab (mAb) Gr-1 (against Ly6c and Ly6g) as previously described (Hickman et al., 2013). Gr-1 treatment eliminated most Ly6c^high^ monocytes in both the blood and PLN.

**Figure 6:**
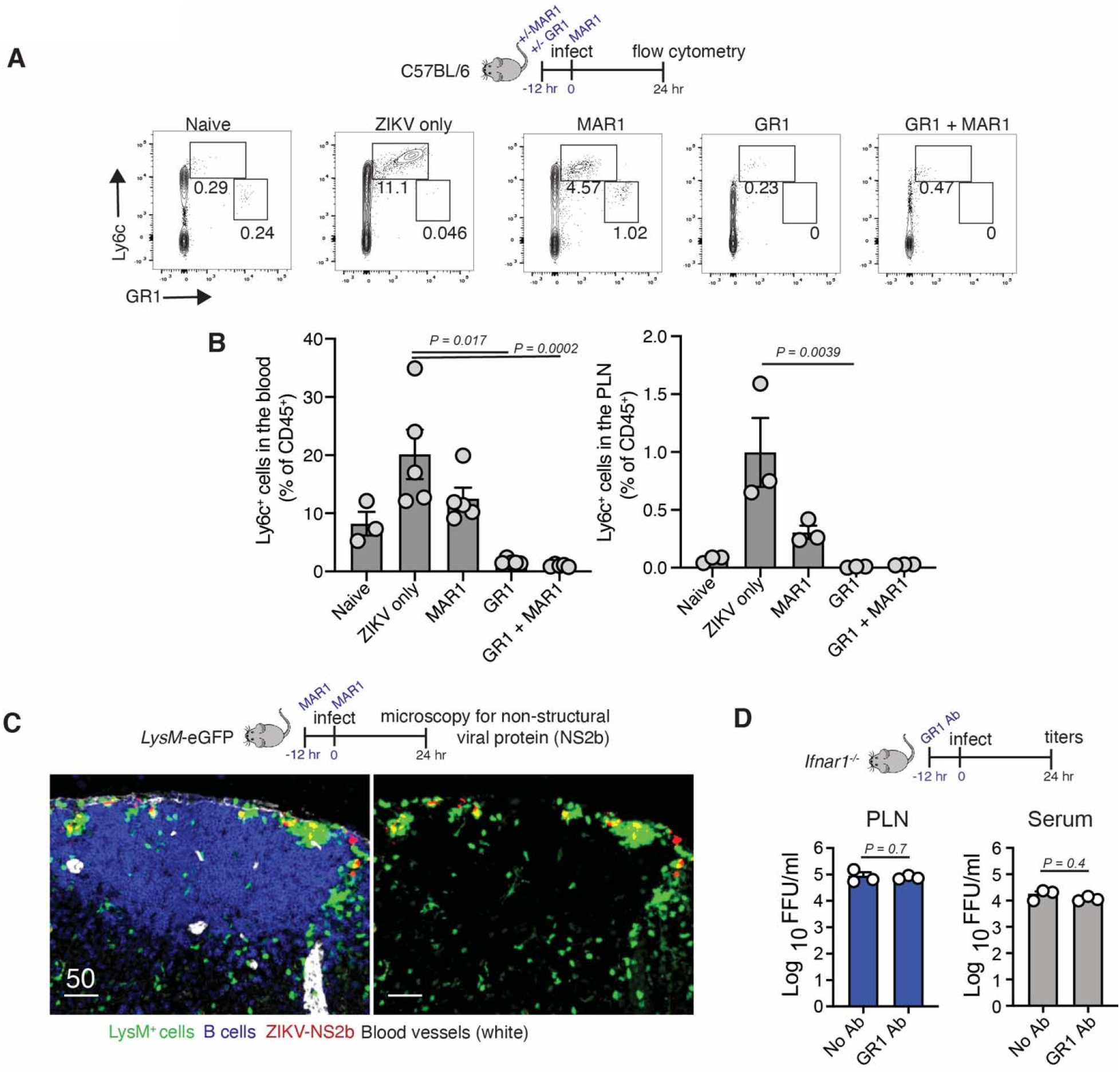
Monocytes are not needed for early ZIKV dissemination. **(A)** Flow plots generated from single cell suspensions of PLNs harvested from naïve C57BL/6 mice (left) or infected C57BL/6 mice at 24 hr p.i. (all other panels). Where indicated, mice were given MAR1 or Gr1 Ab before infection. An additional dose of MAR1 was given at the time of infection. Cells were first gated on CD45^+^ cells. Gating indicates Ly6c^+^ monocytes (top boxes) and Ly6g^+^ neutrophils (middle boxes). **(B)** Frequency of Ly6c^+^ monocytes in the blood (left) or PLN (right) of indicated mice at 24 hr p.i. Statistics = One-way ANOVA. Dots show pooled LNs from individual mice. Scalebars = SEM. Experiment was repeated 2 times with 3-5 mice/group. **(C)** Confocal images of frozen PLN sections from nodes harvested 24 hr p.i. from LysM-eGFP mice treated with MAR1. B220 = blue; LysM-GFP^+^ cells = green; blood vessel (CD31) = white; and ZIKV NS2b protein = red. Right panels omit B cells for clarity. Scalebars =μms. **(D)** Viral titers (FFU/ml) in the PLN (left, blue bars) or serum (right, grey bars) harvested 24 hr p.i. of *Ifnar1*^-/-^ mice with 10^4^ FFU of ZIKV. Mice were given Gr1 Ab prior to infection. Experiment was repeated 3 times with 3-4 mice per group. Dots represent individual mice (either pooled LNs or separate serum) and the average of technical replicates. Dashed line = limit of detection for assay. Statistics = Unpaired t test.

Using confocal and multiphoton microscopy, we examined PLN sections (either live or frozen-fixed) for the presence of inflammatory monocytes in LysM-eGFP reporter mice, which possess green myelomonocytes of varying GFP intensity (neutrophils (bright), monocytes (bright-intermediate), and macrophages (dim)) (Faust et al., 2000). While eGFP-bright neutrophils accumulated in and near the SCS around ZIKV-infected and dying macrophages, we did not detect NS2b-expressing eGFP^+^ cells at the timepoints examined (8-24 hr pi.) (**Figure 6C** and **Supplementary Figure 3)**. Further, depletion of monocytes did not impact infectious viral titers in the PLN or serum at 24 hr p.i. (**Figure 6D**).

Collectively, these data indicate that Ly6C^high^ classical monocytes are not trafficking infectious virus from the skin into the LN nor from the LN to the blood. Further, infection of blood monocytes is not required for high levels of blood-borne virus.

### Nodal macrophage infection alone does not induce systemic disease

Having ruled out other cellular sources for ZIKV dissemination from the LNs, we next examined the overall contribution of nodal macrophage infection to sustained viremia (defined here by virus in the blood), which could be detected in *Ifnar1*^-/-^ mice through at least 5 days p.i. (**Figure 7A**). In contrast, viral titers were just above the limits of detection in CD169 cKO mice on day 3 p.i., and infectious ZIKV could no longer be detected in the serum on day 5 p.i. (**Figure 7B**). Likewise, viral titers in the PLN were very low in CD169 cKO mice on day 3 p.i., but not in *Ifnar1*^-/-^ mice, suggesting that there are secondary targets for ZIKV infection in the LN leading to sustained viral production. Consistent with reduced nodal and serum viral levels, CD169 cKO exhibited no weight loss or mortality after inoculation with the same dose of virus that resulted in marked virus-induced morbidity in *Ifnar1*^-/-^ mice (**Figure 7C**). Collectively, our data indicate the nodal macrophages contribute to viral dissemination from the LN to the blood; however, infection of other cells is needed for virus-induced morbidity.

**Figure 7:**
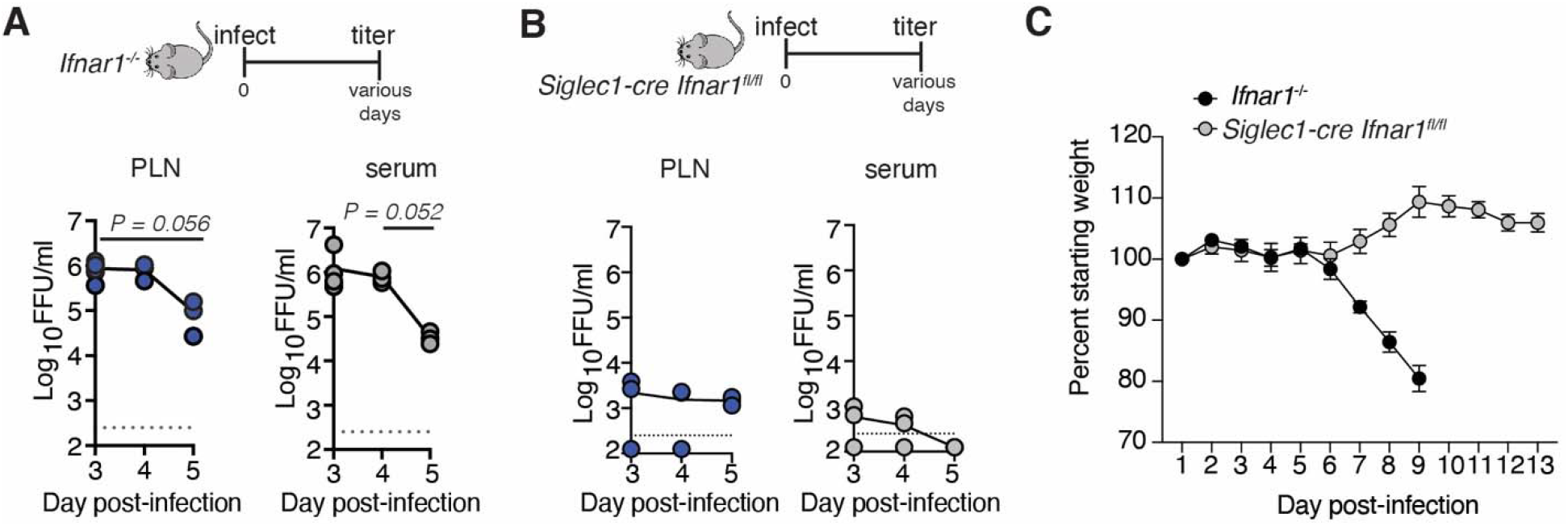
Nodal macrophage infection does not result in morbidity. **(A)** Viral titers (FFU/ml) in the PLN (left, blue dots) or serum (right, grey bars) harvested at the indicated day p.i. of *Ifnar1*^-/-^ mice. Experiment was repeated 2 times with 3-4 mice per group. Results shown are pooled from two independent experiments. Dots represent individual mice (either pooled LNs or separate serum) and the average of technical replicates (values of zero are not shown due to log scale). Dashed line = limit of detection for assay. Values below half the LOD are reported as half the LOD (125 FFU/ml). (**B**) As in (A) but in *Siglec1-cre Ifnar*^fl/fl^ (cKO) mice. **(C)** Weight loss (as a percentage of starting weight) in *Ifnar1*^-/-^ or *Siglec1-cre Ifnar*^fl/fl^ (cKO) mice. Dots show average weight per day p.i. (indicated on x axis). Experiment was repeated twice with 4-5 mice/group. Scalebars = SEM. Statistics = one way-ANOVA.

## Discussion

Many medically significant viral pathogens, including ZIKV and monkeypox virus--both responsible for recent epidemics--follow a similar distribution route through the host over the course of infection. After viral entry via a breach in an epithelial barrier, virus travels to and replicates in the draining LN. After the LN, virus can be detected in the bloodstream and eventually in distal tissues, where the consequences of viral infection are often observable as host pathology. Each step in this process involves the movement of virus into a new part of the body against notable physical and immunological barriers. For example, upon tissue entry, phagocytes of the innate immune system can capture virus to eliminate spread and produce cytokines that limit viral replication in the tissue (Shi and Pamer, 2011). Once virus reaches the blood, endothelial cells form a virus-impermeable primary barrier preventing the free diffusion of virus into the tissue (Mehta and Malik, 2006). While the mechanisms allowing viral circumvention of some of these important controls of systemic homeostasis are well-established, our knowledge of events unfolding in some areas, such as the LN, could be enhanced.

One strategy allowing virus to bypass normal barriers against spread is to hitchhike with immune cells that migrate throughout the body. The number of migratory immune cells, particularly blood monocytes, increases by orders of magnitude during infection. Numerous studies in both mice and humans have identified ZIKV-infected monocytes in the blood or in peripheral organs (Ayala-Nunez *et al*., 2019; Foo *et al*., 2017; Jurado and Iwasaki, 2017; McDonald *et al*., 2020; Michlmayr *et al*., 2017). The results of these studies (along with others during infections with related flaviviruses) have led to the widely accepted viewpoint that myeloid cells are critical for viral dissemination from the skin. However, monocytes could function to spread virus at multiple timepoints during infection. Our studies refine the events leading to ZIKV in the tissues to include LN macrophages as the cells responsible for the initial movement from the LN to the blood. Monocyte involvement primarily must occur later as ZIKV spreads from the blood to distal tissues. Depletion of monocytes did not impact early viral titers in the serum, suggesting that monocytes do not produce this virus but might instead by infected by it. Monocyte infection could occur in either the bloodstream or bone marrow and might be enhanced by increased monocyte mobilization during viremia.

In addition to monocytes, migratory dermal DCs are often implicated in the spread of ZIKV and other flaviviruses from the skin (Begum et al., 2019; Castillo et al., 2019; Duangkhae *et al*.,2018; Hamel *et al*., 2015; King et al., 2020; Lei et al., 2020; McDonald *et al*., 2020; Rathore and St John, 2018; Schneider et al., 2021; Troupin et al., 2016). Given the right stimulation, DCs clearly possess the capacity to migrate from virally infected tissue to the LN (Platt and Randolph, 2013). To our knowledge, however, ZIKV-infected migratory DCs have not been isolated from or identified in the LN. In a previous study, we microscopically examined ZIKV-infected LNs 24 hours after infection and did not identify infected migratory DCs (Reynoso *et al*.,2019). Our data here indicate that ZIKV is present in the blood before large numbers of DCs can migrate into the LN. Future experiments will be required to understand the role of infected DCs at later timepoints. Recent studies with the poxvirus vaccinia (VACV) have demonstrated that the immune system can halt viral spread after cutaneous viral replication by shutting off lymphatic transport of virus or by preventing DC trafficking to the LN (Aggio et al., 2021; Churchill et al., 2022). If these skin immune-defense mechanisms also occur during ZIKV infection, viral replication in the skin might not directly translate to infectious virus in the blood. Nonetheless, local viral replication in the skin is poised to influence systemic viral titers through indirect mechanisms, such as pro-inflammatory cytokine production.

The lymphatics can deliver a first bolus of virus to the LN for capture by strategically positioned macrophages and DCs (Hickman et al., 2008; Junt et al., 2007; Reynoso *et al*., 2019). This early viral delivery initiates the adaptive immune response by providing Ag for B cell activation and allowing for the direct priming of CD8^+^ T cells by LN-resident DCs (Hickman et al., 2011; Junt *et al*., 2007; Reynoso *et al*., 2019). However, given the small number of SSMs in the LN (with some estimates of only 200 SSMs per popliteal LN (Chatziandreou et al., 2017)), we questioned their role as a major source for blood-borne virus. However, our data in CD169 cKO mice suggest that this small group of LN cells can indeed produce enough virus to seed the bloodstream by 12 h post-infection. Future studies will be needed to determine the sources for continued virus output in CD169 cKO mice (for example, other LNs in the chain or CD169^+^macrophages in other tissues, such as the spleen). While high levels of blood-borne virus could still be detected in *Ifnar1*^-/-^ animals at 3 days p.i., viral titers begin to wane in CD169 cKO. This suggests that there is a finite population of these cells that support early but not late ZIKV dissemination.

Subcapsular sinus macrophages undergo attrition during LN infection, vaccination, and inflammation (Gaya *et al*., 2015). The mechanisms leading to SSM death have been debated (e.g. pyroptosis versus necroptosis) (Chatziandreou *et al*., 2017; Gaya *et al*., 2015; Sagoo *et al*., 2016), but SSMs that are not directly infected also die after stimulation. This observation is perplexing as SSM death impairs both LN filtration as well as the development of humoral immune responses to secondary pathogens. Based on our data, we propose that LN macrophage death represents a programmed response to limit virus production by eliminating uninfected cells that could serve as permissive vessels to amplify and disseminate virus.

Collectively, our data reveal the lymphatics and LN sinusoidal macrophages as key participants in the dissemination of ZIKV from the LN to the blood. Early viral spread to the blood occurred in the absence of cellular transport by DCs or monocytes. Together, our data illuminate a timepoint during ZIKV dissemination that could be targeted to prevent downstream infection before viral movement into peripheral tissues.

## Methods

### Mice

*Ifnar1*^-/-^ (Line 314), *Ccr7*^-/-^ (Line 8453), and LysM-eGFP (Line 342) were obtained from the NIAID intramural research repository at Taconic Farms. Wild-type C57BL/6N mice were obtained from Taconic Farms. *Ifnar1^fl/fl^* mice (B6(Cg)-Ifnar1tm1.1Ees/J; #28256) were obtained from Jackson Laboratories. CD169-cre mice have been previously described (Asano et al., 2011). Embryos were obtained from Riken, rederived and bred in-house to generate CD169-cre *Ifnar1^fl/fl^* conditional knockout (cKO) mice. Male and female mice from 6-12 weeks of age were used for experiments. Mice were housed under specific pathogen-free conditions and provided standard rodent chow and sterile water as necessary. All animal studies were approved by and performed in accordance with the Animal Care and Use Committee of the National Institute of Allergy and Infectious Diseases.

### Viral infections and titers

Mice were anesthetized using isoflurane and infected in both hind footpads with 1×10^4^ FFU ZIKV H/PF/2013 unless otherwise indicated (for some experiments, only one footpad was infected). To determine viral titers, LNs or hind feet were collected at various times p.i. and placed in 250 μls of RPMI + 2% FBS + HEPES/LN or 500 μls per foot in metal bead lysing matrix tubes (MP Biomedicals). Samples were homogenized using a Fastprep-24 (MP Biomedicals). Alternatively, blood was collected at the indicated time p.i. into Serum gel Z/1.1 tubes (Sarstedt) and serum separated by centrifugation. To measure ZIKV-induced morbidity, mice were weighed before infection and daily thereafter, and the percent starting weight was calculated for each mouse.

Infectious virus titers were determined using a standard focus forming assay (FFA). Vero cells were seeded at a concentration of 2 x 10^4^ per well in 96-well plates and incubated at 37°C until 90-100% confluency was reached. Confluent monolayers were inoculated in duplicate with LN homogenates or serum and incubated at 37°C for 4 hr. Cells then were overlaid with 1% (wt/vol) methylcellulose (Sigma) in Opti-MEM supplemented (Gibco) with 1% Pen-strep (Gibco) and incubated at 37°C. After 48 hr, cells were fixed with 4%paraformaldehyde for 30 minutes at room temperature. Cells were incubated for 2 hr with ZV-67 mAB in a PBS buffer supplemented with 0.1% saponin and 0.1% BSA. Plates were washed and stained with HRP-conjugated antimouse IgG (Jackson laboratories). Virus-infected foci were visualized using TrueBlue peroxidase substrate (KPL) for 20 min and counted by an ImmunoSpot 5.0.37 microanalyzer (CTL).

### Antibody treatment

MAR1-5A3 Ab was administered intraperitoneally at 1 mg in 1 ml of sterile PBS 12 hr before and the day of footpad infection. For inflammatory monocyte depletion, Gr-1 Ab was administered intraperitoneally at 0.25 mg in 1 ml of PBS 12 hr before infection.

### Flow cytometric analyses after enzymatic tissue dissociation

LNs were collected at various times p.i. and single-cell suspensions prepared by digestion with Liberase (Roche) + DNAse (Worthington) for ~ 1 hr at 37°C. Cells were disrupted by 3 rounds of vigorous pipetting, suspensions were filtered through 60 μm nylon-filter capped FACS tubes. LN cells were stained for 20 min on ice with CD45 (clone 30-F11), CD3 (clone 17A2), B220 (clone RA3-6B2), CD11c (clone N418), CD11b (clone M1/70, BioLegend), CD169 (clone 3D6.112), F4/80 (clone BM8), Gr-1(clone RB6-8C5) and/or (HK1.4) and fixable viability dye Ghost Dye UV450 or Zombie Aqua. Samples were washed twice post-staining in Hanks Balanced Salt Solution + 0.1% BSA and fixed for 20 min with 3.2% paraformaldehyde. Samples were washed twice in PBS and acquired on a Fortessa flow cytometer (BD Biosciences). Flow data were analyzed using FlowJo Software (TreeStar).

### Confocal microscopy of frozen LN sections

Confocal microscopy was performed as described previously (Reynoso *et al*., 2019). Briefly, LNs were removed at the indicated time p.i., fixed in periodate-lysine-paraformaldehyde (PLP) for 48 hr, and moved to 30% sucrose/PBS solution for 24 hr. Tissues were embedded in optimal-cutting-temperature (OCT) medium (Electron Microscopy Sciences), carefully oriented to cut through both the B cell follicles and the medullary sinuses, and frozen in dry-ice-cooled isopentane. 16-μm sections were cut on a Leica cryostat (Leica Microsystems). Sections were blocked with 5% goat, donkey, bovine, rat, or rabbit serum and then stained with one or more of the following Abs: ZIKV NS2b protein Ab (polyclonal, Genetex), ZIKV E protein Ab (polyclonal, Genetex), ERTR7 (rat monoclonal, Abcam), B220 (clone RA3-6B2, Thermo Fisher Scientific), CD169 (clone 3D6.112, BioLegend), Lyve-1 (clone ALY7, Thermo Fisher Scientific), CD11c (clone N418, Thermo Fisher Scientific), SIGN-R1 (clone eBio22D1, Thermo Fisher Scientific), CD31 (clone MEC13.3, BioLegend). Sections were incubated with secondary antibodies as needed and as controls, and images were acquired using identical PMT (photomultiplier tube) and laser settings. Images were processed and analyzed using Imaris software (Oxford Instruments).

### ZIKV reporter virus particle production and neutralization assay

Reporter virus particles (RVPs) incorporating the structural proteins of ZIKV (strain H/PF/2013) were produced by complementation of a sub-genomic GFP-expressing replicon derived from a lineage II strain of WNV as previously described (Dowd et al., 2016a). Briefly, HEK-293T cells were transfected with plasmids encoding the WNV replicon and ZIKV structural genes at a 1:3 ratio by mass using Lipofectamine 3000 (Invitrogen), followed by incubation at 30°C. RVP-containing supernatant was harvested from cells at days 3-6 post-transfection, filtered through a 0.22 μm filter, and stored at −80°C. To determine virus titer, two-fold dilutions of RVPs were used to infect Raji cells that express the flavivirus attachment factor DC-SIGNR (Raji-DCSIGNR) in duplicate technical replicates at 37°C. GFP-positive infected cells were detected by flow cytometry 2 days later. In subsequent neutralization assays, RVPs were sufficiently diluted to within the linear range of the virus-infectivity dose-response curve to ensure antibody excess at informative points. For neutralization studies, ZIKV RVPs were mixed with serial dilutions of heat-inactivated mouse sera for 1 h at 37°C, followed by infection of Raji-DCSIGNR cells in duplicate technical replicates. Infections were carried out at 37°C and GFP-positive infected cells quantified by flow cytometry 48 hr later. Results were analyzed by non-linear regression analysis to estimate the dilution of sera required to inhibit 50% of infection (EC_50_). Samples were initially tested at a starting dilution of 1:60 (based on the final volume of cells, virus, and sera per well), which was designated as the limit of detection; neutralization titers predicted by non-linear regression as <60 were reported as a titer of 30 (half the limit of detection). The reported EC50 values represent the average of 2 independent neutralization assays.

### Statistical analyses

Significances were calculated using Prism V 8.3.0 (Graphpad Software) using unpaired twotailed Mann-Whitney t-tests (when only two groups were present) or using a one-way ANOVA with multiple comparisons as indicated in the Figure legends.

## Acknowledgments

This work was supported by the Intramural Research Program of the NIAID, NIH. The graphical abstract was drawn by Ethan Tyler, NIH Medical Arts Branch.

## Author Contributions

Conceptualization, H.D.H.; methodology, H.D.H., G.V.R, D.G. and T.C.P; materials, M.T.; investigation, G.V.R., D.G., A.K., C.C.A, M.A., J.P.S., S.M.V., C.R.C., C.S.M., K.A.D, S.M., and H.D.H.; writing-original draft, H.D.H.; writing-review and editing, all authors; supervision, H.D.H.,T.C.P; funding acquisition, H.D.H., T.C.P.

## Declaration of Interests

The authors declare no competing interests.

## Notes

### Competing Interest Statement

The authors have declared no competing interest.

