## supplemental figures 1-3 for "Zika virus spreads through infection of lymph node-resident macrophages"

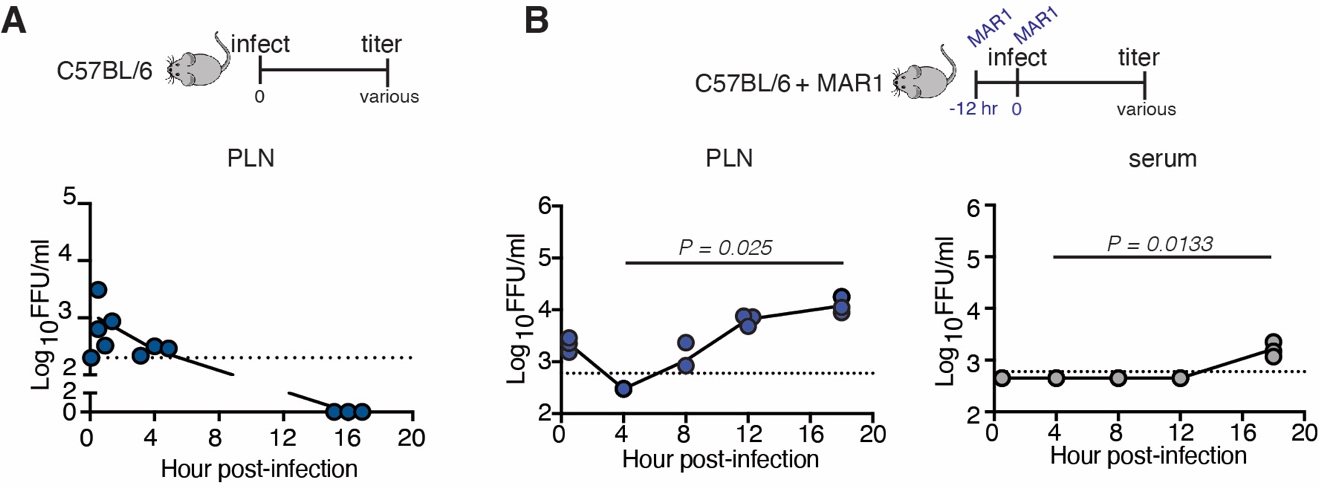

**Supplemental Figure 1: ZIKV replicates in the popliteal LN.**

**(A)** Viral titers (in focus-forming units (FFU)/ml)) in the PLN harvested at the indicated time (in hrs) following footpad (FP) injection of C57BL/6 mice with 10^4^ FFU of ZIKV H/PF/2013. Dots represent individual mice (pooled PLNs) and the average of technical replicates in the focus-forming assay (FFA). A dashed line shows the limit of detection of the FFA. **(B)** Viral titers in the PLN (left, blue dots) or serum (right, grey dots) at the indicated timepoint during the first 20 hr p.i. of C57BL/6 mice treated with the anti-Ifnar1-Ab MAR1-5A3.

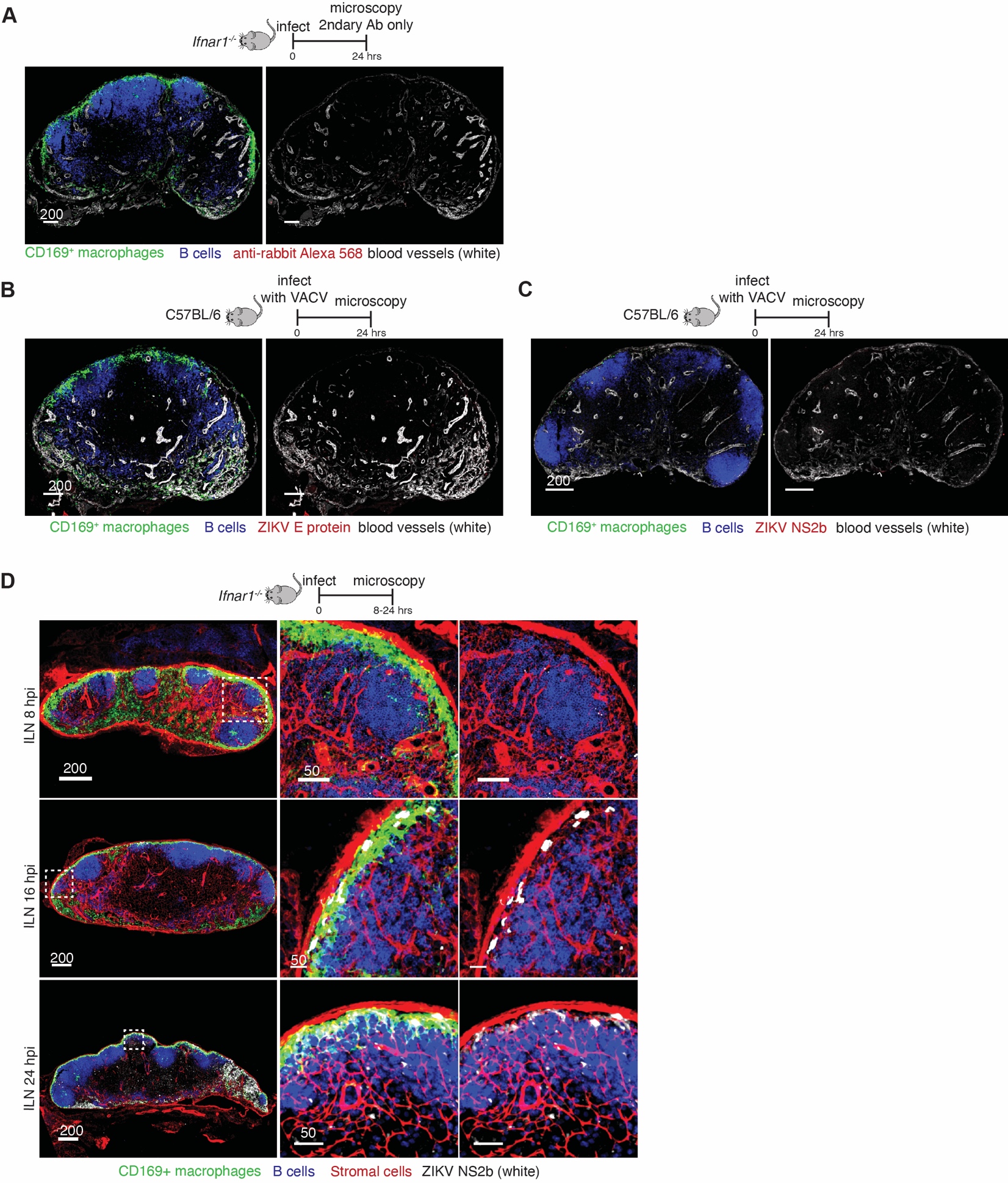

**Supplemental Figure 2: ZIKV infects SSMs in the iliac LN.**

**(A)** Confocal images of frozen LN sections from PLNs harvested 24 hr p.i. from *Ifnar1*^-/-^ mice infected with ZIKV. Nodes were stained with secondary Ab only to reveal background levels of Ab binding. Left panel shows all stains. Right panel omits CD169 and B220 staining to reveal secondary Ab staining more clearly. Green = CD169^+^ macrophages; red = secondary Ab; white = blood vessels; blue = B cells. **(B)** Confocal images of frozen LN sections from PLNs harvested 24 hr p.i. from C57BL/6 mice infected with vaccinia virus (VACV). Left panel shows all stains. Right panel omits CD169 and B220 staining to reveal secondary Ab staining more clearly. Green = CD169^+^ macrophages; red = ZIKV E protein; white = blood vessels; blue = B cells. **(C)** As in (B) but with ZIKV NS2b primary Ab. Green = CD169^+^ macrophages; red = ZIKV NS2b protein; white = blood vessels; blue = B cells. **(D)** Confocal images of frozen LN sections from iliac LNs harvested at the indicated time (shown in hours p.i.) from *Ifnar1*^-/-^ mice. Left panels show the entire ILN. Middle panel shows a higher magnification image of SSMs. Right panel omits CD169 staining to more clearly reveal NS2b staining. Green = CD169^+^ macrophages; red = LN stroma; white =ZIKV NS2b protein; blue = B cells. Images are representative of 6-10 LNs/timepoint harvested from 3-5 mice. Scalebars = µms.

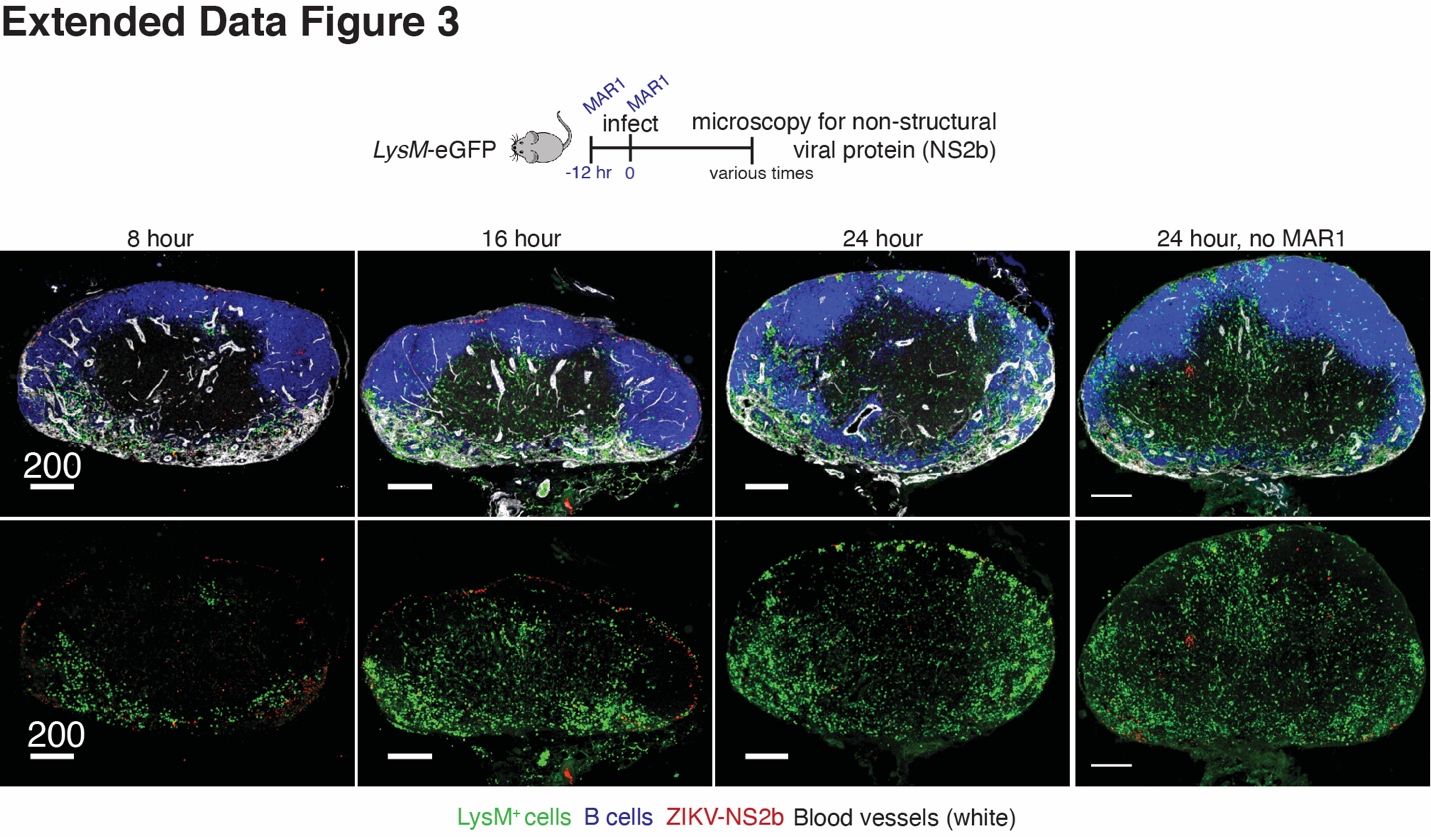

**Supplemental Figure 3: Myelomonocytes accumulate around ZIKV-infected LN macrophages.**

Confocal images of frozen PLN sections from nodes harvested at the indicated hr p.i. from LysM-eGFP mice treated with MAR1. B220 = blue; LysM-GFP^+^ cells = green; blood vessel (CD31) = white; and ZIKV NS2b protein = red. Bottom panels omit B cell and blood vessel staining for for clarity. Scalebars = µms.
